# Learning Cortical Hierarchies with Temporal Hebbian Updates

**DOI:** 10.1101/2023.01.02.522459

**Authors:** Pau Vilimelis Aceituno, Matilde Tristany Farinha, Reinhard Loidl, Benjamin F. Grewe

## Abstract

A key driver of mammalian intelligence is the ability to represent incoming sensory information across multiple abstraction levels. For example, in the visual ventral stream, incoming signals are first represented as low-level edge filters and then transformed into high-level object representations. These same hierarchical structures routinely emerge in artificial neural networks (ANNs) trained for image/object recognition tasks, suggesting that a similar process might underlie biological neural networks. However, the classical ANN training algorithm, backpropagation, is considered biologically implausible, and thus several alternative biologically plausible methods have been developed. For instance, several cortical-inspired ANNs in which the apical dendrite of a pyramidal neuron encodes top-down prediction signals have been proposed. In this case, akin to theories of predictive coding, a prediction error can be calculated locally inside each neuron for updating its incoming weights. Notwithstanding, from a neuroscience perspective, it is unclear whether neurons could compare their apical vs. somatic spiking activities to compute prediction errors. Here, we propose a solution to this problem by adapting the framework of the apical-somatic prediction error to the temporal domain. In particular, we show that if the apical feedback signal changes the postsynaptic firing rate, we can use differential Hebbian updates, a rate-based version of the classical spiking time-dependent plasticity (STDP) updates. To the best of our knowledge, this is the first time a cortical-like deep ANN has been trained using such time-based learning rules. Overall, our work removes a key requirement of biologically plausible models for deep learning that does not align with plasticity rules observed in biology and proposes a learning mechanism that would explain how the timing of neuronal activity can allow supervised hierarchical learning.

## 1 INTRODUCTION

To survive in complex natural environments, humans and animals transform sensory input into neuronal signals which in turn generate and modulate behavior. Learning of such transformations often amounts to a non-trivial problem, since sensory inputs can be very high-dimensional and complex. To form hierarchies, cortical networks need to process these sensory signals and convey plasticity signals down to every neuron in the hierarchy so that the output of the network (e.g., the motor output or behavior) improves during learning. In deep learning, this is known as the credit assignment (CA) problem and it is commonly addressed by the error backpropagation (BP) method. During BP learning, neurons in the lower hierarchies change their afferent synapses by integrating a backpropagated error signal. A neuron’s afferent weight update is then calculated as the product of the presynaptic activity and its non-local output error. However, several key aspects of BP are still at odds with learning in biological neural networks (Crick, 1989; Lillicrap et al., 2020). For example, ANNs separate the processing or encoding of neuronal activity signals from the weight update signals, they utilize distinct phases and they implement an exact weight symmetry of forward and feedback pathways. Moreover, plasticity in biological synapses is local in space and time and tightly coupled to the timing of the pre- and post-synaptic activity (Bi and Poo, 1998).

Attempting to address some of these implausibilities, recent cortical-inspired ANN models leverage network dynamics to directly couple changes in neuronal activity to weight updates (Whittington and Bogacz, 2017; Sacramento et al., 2018). Those models postulate multi-compartment pyramidal neurons with a highly specialized dendritic morphology that use their apical dendrite to integrate a feedback signal that modulates feedforward plasticity (Fig. 1A, left schematic). Although multi-compartment models agree with some biological constraints, such as the spatial locality of learning rules and the fact that feedback not only generates plasticity but also affects neuronal activity (Gilbert and Li, 2013), the apical ‘dendritic-error’ learning approach still requires tightly coordinated and highly specific error signaling circuits.

**Figure 1.**
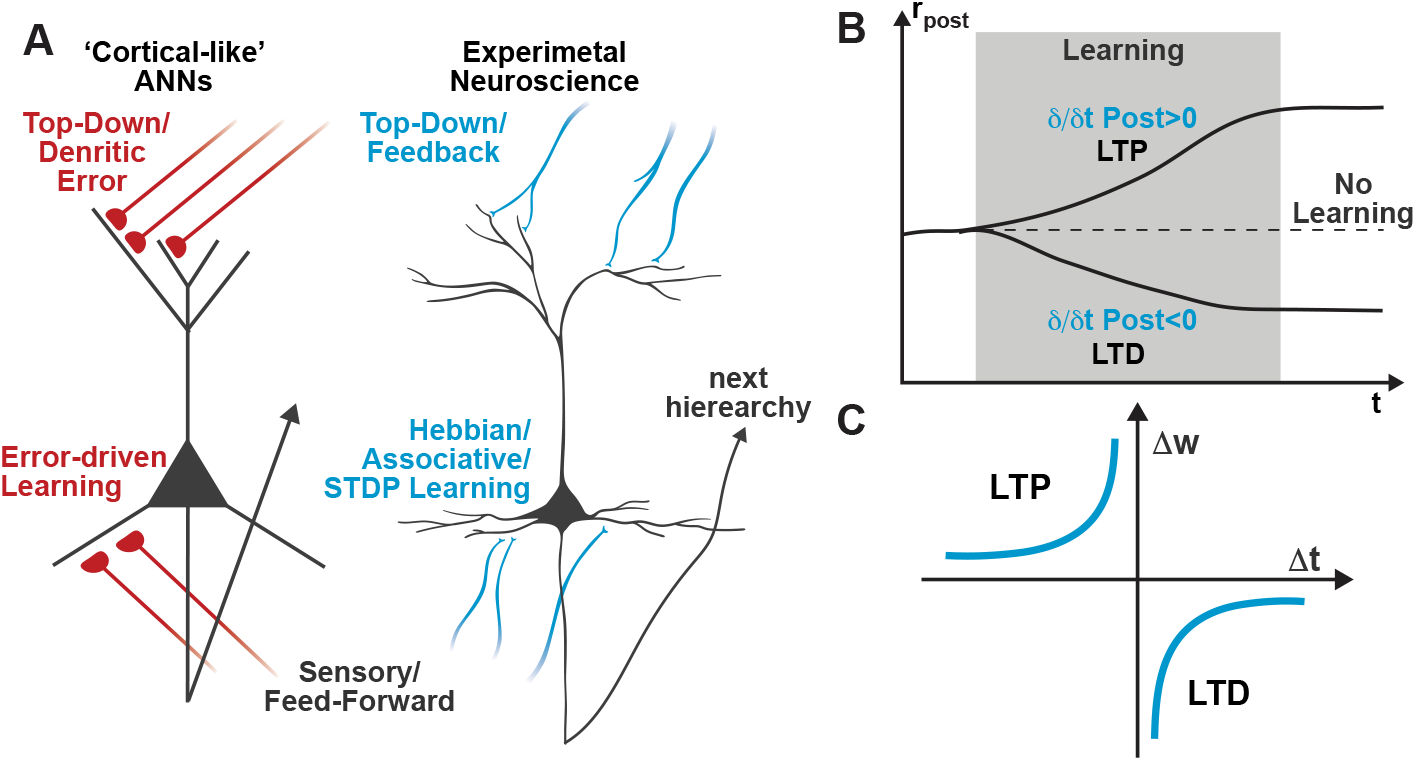
Schematic comparison of learning rules in artificial and biological neural networks. (A) While recently proposed cortical-like ANNs utilize dendritic-error learning rules to induce plasticity in basal synapses (left neuron), biologically observed plasticity rules are based on Hebbian-type associative learning rules such as STDP (right neuron). (B) A temporal Hebbian update rule such as STDP directly relates to increasing or decreasing postsynaptic activity. Thus, STDP learning is also often referred to as differential Hebbian learning Xie and Seung (1999); Zappacosta et al. (2018). (C) Classical STDP profile showing ranges of Δt that induce long-term potentiation (LTP) and long-term depression (LTD), as extracted from experimental observations in neuroscience.

To tackle these issues, we recently developed a novel class of cortical-inspired ANNs that utilize the same dendritic-error learning rule, but do not require highly specific error circuits and are capable of online learning of all weights without requiring separate backward and forward passes (Meulemans et al., 2021a,b, 2022). In this model, also known as ‘Deep Feedback Control’ (DFC), we dynamically tailored the feedback to each hidden neuron until the network output reaches the desired target. The weight update of the feedforward pathway is then calculated upon convergence as the difference in neural activities when fully taking into account the effect of top-down apical feedback. Still, this model relies on the same dendritic-error learning rule as its predecessors (Whittington and Bogacz, 2017; Sacramento et al., 2018), and it is unclear how a neuron would be able to compare the activities of its basal and apical compartments (Fig. 1A, left scheme). In this work, we argue that compartment-based dendritic learning rules can be substituted by experimentally validated temporal Hebbian learning rules (e.g., STDP).

The gist of our argument is that single-compartment neurons, whose firing rate is strongly affected by apical input, can use the difference between consecutive instances of their activity as a learning signal (Fig. 1B), as opposed to comparing the changes in two different compartments. Based on this dynamic change in the postsynaptic activity we can thus encode the learning signal while being consistent with experimentally observed learning rules – such as STDP (Fig. 1C).

In our framework, the constraints imposed on the learning dynamics and the architecture agree with experimental evidence beyond the temporal-based Hebbian learning, making our work relevant for experimental neuroscientists (discussed in Section 3). As a first step, we show in the next section that for a single neuron the weight updates obtained by differential Hebbian (DH), as well as by the STDP, can be seen as equivalent to the existing rules of alternative cortical-inspired as well as standard BP networks (Bengio et al., 2017; Xie and Seung, 2003).

Spike-Timing Dependent Plasticity (STDP)

When using the term STDP, we here refer to the well-established observation that the precise timing of pre- and post-synaptic spikes significantly determines the sign and magnitude of synaptic plasticity (Markram et al., 1997; Bi and Poo, 1998). In cortical pyramidal neurons, a presynaptic spike that precedes a postsynaptic spike within a narrow time window induces long-term potentiation (LTP) (Markram et al., 1997; Bi and Poo, 1998; Feldman, 2012; Nishiyama et al., 2000; Sjöström et al., 2001; Wittenberg and Wang, 2006); if the order is reversed it leads to long-term depression (LTD). Using this classical STDP profile (Fig. 1A), multiple theoretical models were able to predict biological plasticity by assuming a simple superposition of spike pairs (Gerstner et al., 1996a; Kempter et al., 1999; Abbott and Nelson, 2000; Song et al., 2000; van Rossum et al., 2000; Gütig, 2016; Izhikevich and Desai, 2003).

## 2 RESULTS

### 2.1 Single neuron supervised learning with STDP

We first demonstrate how an STDP learning rule can be used to train a single neuron on a linear classification task (Fig. 2A). We use a neuron with the sigmoid activation function, which gets both feedforward basal inputs from two other neurons (A and B) and a feedback apical input, resulting in the rate-based dynamics

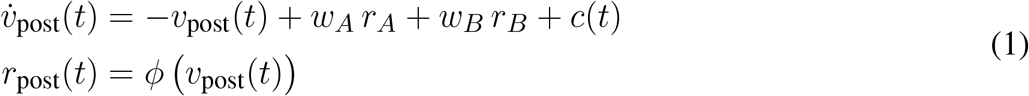

**Figure 2.**
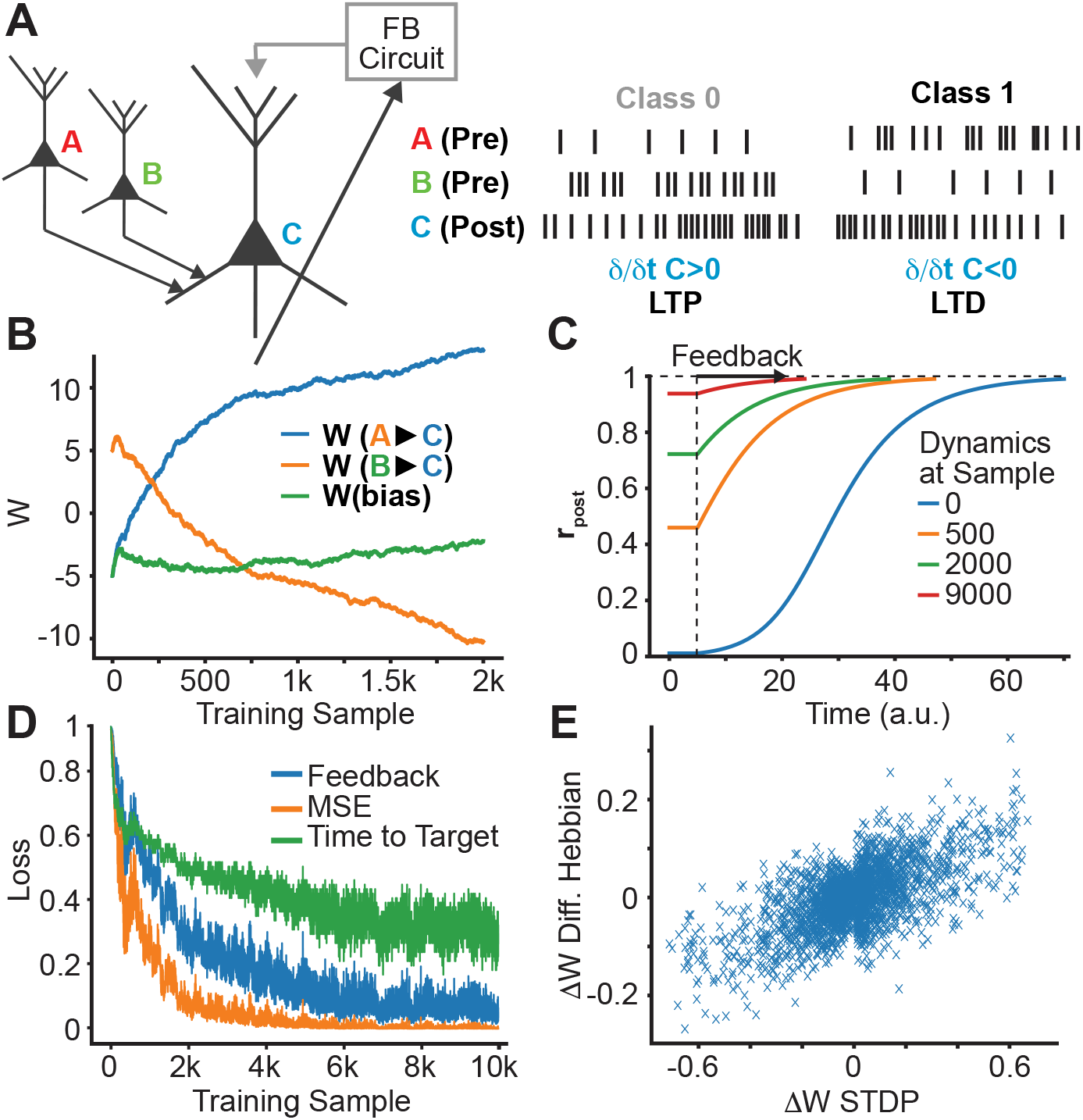
Single neuron supervision and STDP learning. (A) Single neuron supervision scheme. Throughout learning, we plot: (B) the evolution of the presynaptic weights originating from neurons A and B; (C) the evolution of the neural dynamics; (D) the decreasing feedback, MSE loss, and time-to-target; (E) the correlation between DH and STDP weight updates.

where *w*_*A*_, *w*_*B*_ are the synaptic strengths of the connection from the input neurons to the output neuron, *r*_*A*_, *r*_*B*_ are the firing rates of neurons *A, B*, respectively, *r*_post_(*t*) is the output firing rate at time *t*, and *c*(*t*) is the apical feedback given to the output neuron. In our simple example, the neuron can get two incoming stimuli, from neurons *A* and *B*, and the apical feedback *c*(*t*) changes the output firing rate to be high when *B* is presented and low when *A* is presented.

The firing rate variables are converted into spike trains with an inhomogeneous Poisson process, where at every time step the probability of spiking in each neuron is given by *r*_post_(*t*), *r*_*A*_, *r*_*B*_, respectively. These spike trains are then used to induce synaptic weight changes by STDP (see Fig. 2B).

We observe in Figure 2B that, as learning progresses, the weights evolve to the expected values (high for *w*_*B*_, low for *w*_*A*_), and that this changes the dynamics of *r*_post_(*t*), causing *r*_post_ to start closer to its target value and, thus, shortening the time and the feedback required to produce the desired output (Fig. 2C,D). Such changes can be understood in terms of the following set of equivalent loss functions that are minimized:

- The initial distance to the target activity can be computed as the Mean Squared Error (MSE), denoted by ℒ. This loss is commonly used in the machine learning literature as a standard measure of performance.
- The feedback required to maintain or reach the target activity, denoted by ℋ. This loss is equivalent to the one presented in previous works on using feedback to train neural networks (Meulemans et al., 2020; Gilra and Gerstner, 2017) and relates to the intuition from predictive coding that a trained ANN minimizes the feedback needed to correctly process the input (Rao and Ballard, 1999).
- The time delay to reach the target is denoted by 𝒯. This loss function represents the amount of time a neuron takes to reach its target value. This idea appears in previous works based on STDP models (Masquelier et al., 2009; Vilimelis Aceituno et al., 2020) and is also implicitly used in models for learning in deep networks (Luczak et al., 2022a).

To relate the three losses to temporal Hebbian learning, we re-express the STDP update through its rate-based form, known as the differential Hebbian (DH) learning rule (Xie and Seung, 1999; Saudargiene et al., 2004; Bengio et al., 2017),

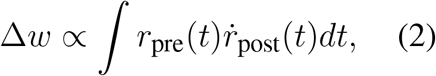

where Δ*w* is the change in feedforward synaptic strength, *r*_pre_(*t*) is the presynaptic activity and 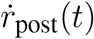 is the derivative of the postsynaptic activity, which corresponds to the change in firing probability. As we see in Figure 2E, the DH learning rule is indeed similar to STDP, albeit with noise induced by the inherent stochasticity of using a Poisson neuron.

To understand how this rule relates to the loss functions mentioned above, we note that in the single neuron example the presynaptic firing rate is fixed, which simplifies the previous rule to

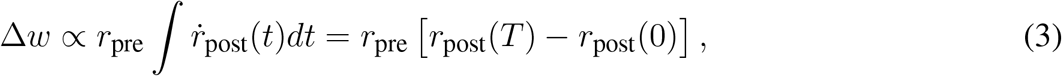

where *r*_post_(*T*) is the postsynaptic activity after reaching the target state. This corresponds to the dendritic-error learning rule (Sacramento et al., 2018; Meulemans et al., 2020; Gilra and Gerstner, 2017). The correlation between this rule and STDP is shown in Figure 2E. In this single neuron setting, it is clear that both STDP and DH learning decrease the three loss functions: having an initial activity that is closer to the target activity implies that the MSE loss is lower at the beginning, and also that the change in activity is smaller, hence it needs less feedback and the target can be reached much faster (see Appendix I for a detailed explanation). The next key question is whether STDP and DH learning can be adopted for training multilayer neural networks in the same manner.

### 2.2 Learning multilayer networks with DH

In order to extend our results to multilayer neural networks, each neuron must receive top-down feedback signals. To compute the appropriate feedback signals, we use the framework of deep feedback control (DFC) from our previous work (Meulemans et al., 2021a). In DFC, the neurons receive a feedback signal that is computed by a controller whose goal is to achieve a target output response (Fig. 3A). This controller can be implemented by a standard PI controller which feeds back into each neuron with a weight that can be either calculated analytically or learned by a DH learning rule. In previous work, we showed that DFC can train deep neural networks to attain state-of-the-art performances, but only when combined with a dendritic (two-compartment-based) learning rule.

**Figure 3.**
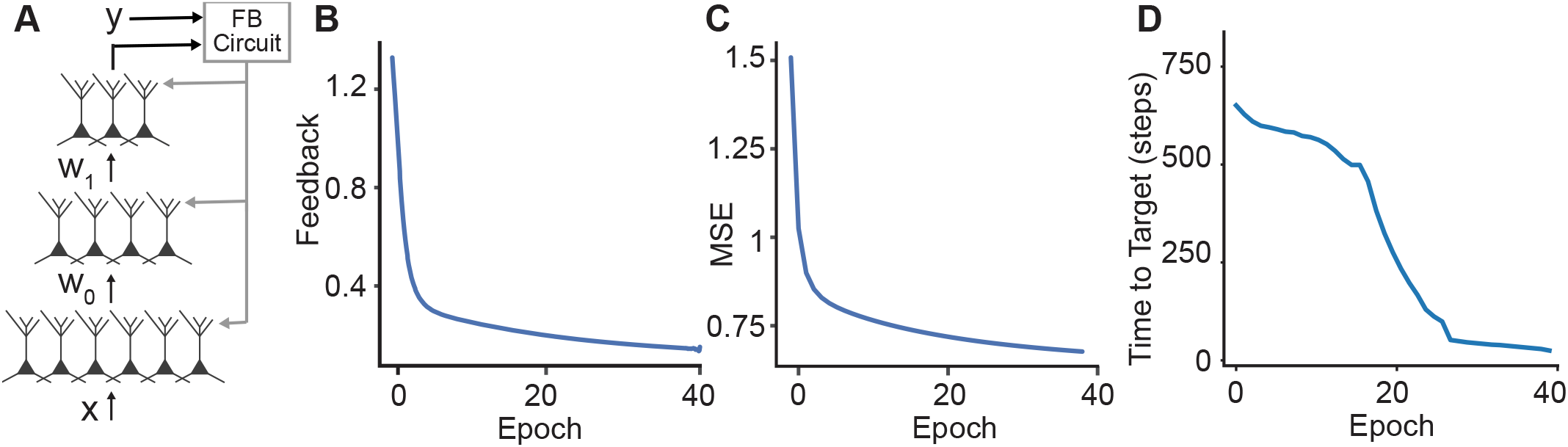
Learning in hierarchical networks. (A) Schematic illustration of how the feedback can be used to train a deep network. We use the desired output label and the output of the network to compute the feedback signals that are sent to the apical dendrites of the neurons. (B-D) Throughout the training, the three losses (feedback, MSE loss, and time-to-target) decrease for the MNIST benchmark.

However, in contrast to the single neuron setting, the DH learning rule in a multiplayer network is not equivalent to the dendritic-error learning rule. Since the presynaptic firing rate of most synapses changes in time, the DH learning rule can then be expressed as

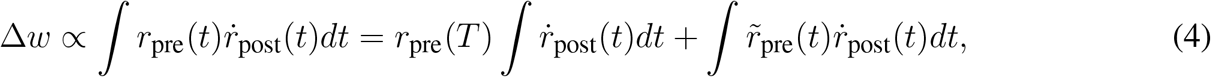

where the extra term includes 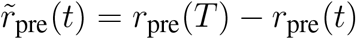 which is the difference between the presynaptic firing rate at time *t* and its target.

To understand why DH learning works despite being different from the classical dendritic-error learning rule, it is useful to note that the term 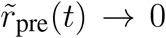 as learning proceeds. This can be explained by the following inductive argument. For the first layer, the presynaptic input is static, so 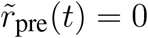. The synapses between the first and second layers will adapt and eventually converge to values that make the neurons in the second layer produce the target firing rate without needing feedback at all. Therefore, the presynaptic firing rate will become static and, thus, 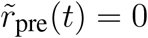 for the second layer. Then, the second layer will receive a static presynaptic input, allowing us to apply this reasoning across the network. This implies that the fixed points of the synaptic weights (the weight configurations where Δ*w* = 0) with dendritic-error learning are the same as the fixed points with DH learning (for each training example). A more detailed derivation is presented in Appendix II.1.

Knowing that the learning dynamics of DH have the same attractive fixed points as those of the dendritic-error learning used in DFC and that those of DFC are in turn equivalent to those in BP (Meulemans et al., 2021a), we conclude that these three training methods similarly train a deep neural network.

We test this on the MNIST computer vision benchmark and show that DH with feedback computed through the DFC framework can train a three-hidden layer network (256×256×256) to match state-of-the-art performances. We compare our framework with BP and the original DFC framework based on dendritic-error learning in Table 1. For the MNIST benchmark, we find that the performances of BP, DFC, and DH-DFC are comparable.

**Table 1.** MNIST Performance for DH-DFC, DFC, and BP. We compare the accuracies reached by our training procedure (DH-DFC) with those achieved by BP and DFC (as reported in Meulemans et al. (2021b)) in the MNIST dataset, achieving competitive performances.

|  | MNIST |
| --- | --- |
| BP | $1.74 \pm 0.10\%$ |
| DFC | $1.98 \pm 0.05\%$ |
| DH-DFC | $1.89 \pm 0.15\%$ |

To complement our analysis, we investigate the training loss in DH-DFC. We note that the amount of feedback required to reach the target decreases throughout the training (Fig. 3B), implying that DH-DFC also decreases the required feedback. The MSE loss also decreases (Fig. 3C), hence DH-DFC also learns by minimizing an implicit error. Finally, we show that the latency to reach the target is also reduced (see Fig. 3D), implying that the latency-reduction nature of temporally asymmetric learning rules (Masquelier et al., 2009; Vilimelis Aceituno et al., 2020) is reflected in our framework.

We also evaluate the similarity of the weight updates given by different learning rules in Figure 4. We find that both the DFC and the BP updates are strongly positively correlated with DH-DFC, with coefficients of determination of 0.804 and 0.966, respectively.

**Figure 4.**
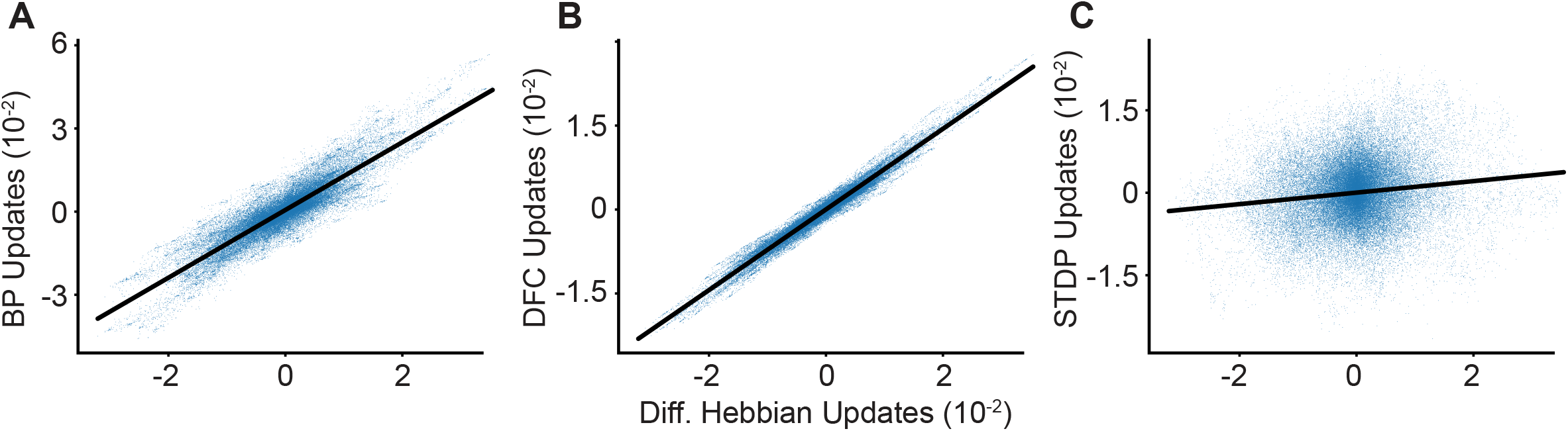
Comparison of learning algorithms. We calculate the synaptic weight update in our deep network using different algorithms and synaptic plasticity rules. We compare our DH weight updates (x-axis) to the updates given by other algorithms. Weight update correlations between DH-DFC and: (A) BP; (B) DFC; (C) and STDP. We observe a clear correlation between DH-DFC and both BP and DFC, and a significant but weaker correlation with STDP.

Finally, we compare the DH-DFC weight updates with STDP updates, which have a positive correlation, with a coefficient of determination of 0.008, but a very noisy alignment due to the randomness induced by using Poisson neurons (see Appendix III). This randomness can be reduced by computing several parallel conversions of firing rates to Poisson spike trains and averaging the resulting STDP updates, although here we find that limitations in computer memory prevent us from reaching state-of-the-art accuracies (see Appendix III).

## 3 DISCUSSION

Building upon previous studies, our work represents an important step forward to explore the different aspects of hierarchical learning in biological networks, including phenomenological and physiological learning rules, bioplausible deep learning models, and predictive coding. In this section, we discuss how our work relates to these aspects.

### 3.1 How does our work fit into the existing literature

#### 3.1.1 Learning with time-dependent hebbian learning rules

Temporal Hebbian learning rules such as STDP or DH rules have been mostly used for unsupervised learning (Sjöströ m et al., 2010; Lazar et al., 2009; Toyoizumi et al., 2005; Gerstner et al., 1996b) or as an enhancement of supervised learning in shallow networks (Diehl and Cook, 2015). In order to use these rules in a supervised setting, it is required to have a teaching signal, which can be implemented either through a neuromodulator or a third-factor learning rule (Frémaux and Gerstner, 2016). However, such approaches do not go beyond shallow networks (Illing et al., 2019). Although it has been suggested that STDP or DH could be adopted for error-driven hierarchical learning (Hinton et al., 2007; Xie and Seung, 1999; Bengio et al., 2017), a suitable network architecture and dynamics to combine time-dependent Hebbian learning rules with deep networks has not been proposed yet (Bengio et al., 2015). Our work fills this gap by providing a method that is able to train deep hierarchies with a learning rule that retains the time-based principles of STDP.

Furthermore, in theoretical neuroscience, causal Hebbian learning rules such as STDP render neurons to fire earlier in time with respect to the onset of an incoming stimulus (Masquelier et al., 2009; Vilimelis Aceituno et al., 2020). This is captured in our model, where neurons change to reach their future (target) activity progressively earlier, thus minimizing the time to target. We note that having neurons in a deep neural network that can learn by predicting their own neuronal activity is known in the deep learning literature (Luczak et al., 2022b), but has not been explored with experimentally validated biological learning rules.

#### 3.1.2 Predictive coding and top-down feedback

Due to the close relation of our work to predictive coding (PC), we next explain in detail the similarities and differences of our approach to PC. In the PC literature, learning decreases the amount of feedback that goes from upper to lower layers, and this process intrinsically generates expedited neuronal responses after stimulus presentation, which are often interpreted as predictions (Whittington and Bogacz, 2017; Friston and Kiebel, 2009). The PC framework goes beyond explanations of these activities by proposing neural circuits that could implement this behavior (Rao and Ballard, 1999; Bastos et al., 2012). However, predictive coding as a mechanistic theory for neural circuits requires explicit error encoding (Rao and Ballard, 1999; Bastos et al., 2012; Koch and Poggio, 1999), a requirement which is problematic for making valid testable predictions (Kogo and Trengove, 2015). In contrast, our framework can exhibit a similar reduction of top-down feedback and anticipated neuronal responses, but since it is based on the target activities of neurons rather than on errors, it does not require explicit errors to be encoded. This shows that it is possible to design neural circuits that can reproduce the relevant PC features where errors are not explicitly encoded but rather represented implicitly by the temporal neuronal dynamics. To illustrate this effect, in Figure 5 we plot the feedback that modulates a deep network during training. The feedback decreases as the model learns but, when we randomly shuffle the labels – which can be considered as a surprising response – the feedback signal increases substantially, thereby changing the neuronal activity.

**Figure 5.**
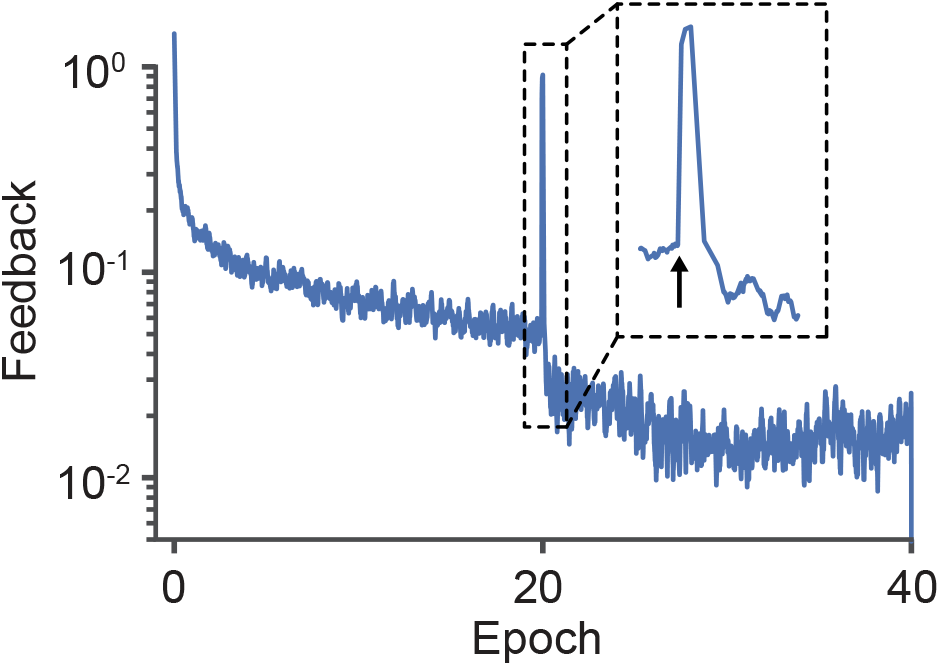
Surprise triggers a large feedback signal that alters neuronal activities. Across learning, the change in the postsynaptic activity driven by the apical feedback reduces as neurons reach their target rates. However, when the labels are randomly swapped (indicated by the arrow in epoch 20), the apical feedback is notably increased. Note that the label switch did not set the network to its baseline state, because the required feedback decreased to the pre-shift level much faster than on the first epochs.

#### 3.1.3 Alternative bioplausible deep learning models

Other bioplausible deep network models such as equilibrium propagation, dendritic-error learning, or burst propagation require a learning signal to be computed either by using two separate phases (Scellier and Bengio, 2017), distinct dendritic and somatic compartments (Sacramento et al., 2018) or via multiplexing of feedback and feedforward signals as bursts and single spikes (Payeur et al., 2021), respectively. In contrast, our model encodes supervision signals as temporal changes in postsynaptic activities, which arrive at individual neurons via their apical dendrite with a short time delay. Table 2 provides a comprehensive comparison of our approach to the most recent alternative bioplausible deep learning methods and how they relate to experimental observations.

**Table 2.** Comparison of diverse bioplausible hierarchical learning methods. For further details on these methods: Equilibrium Propagation (Scellier and Bengio, 2017), Predictive Coding (Whittington and Bogacz, 2017), Dendritic-Error (Sacramento et al., 2018), Burst Propagation (Payeur et al., 2021). Note that the MNIST performance comparison does not take into account different model sizes.

| Method | Implicit Error |  | Explicit Error |  |  |
| --- | --- | --- | --- | --- | --- |
|  | Temporal Hebbian | Equilibrium Propagation | Predictive Coding | Dendritic-Error | Burst Propagation |
| Error Type | Control | Contrastive | Prediction | Prediction | Multiplexed |
| Phases / Compartments | 1/1 | 2/1 | 1/2 | 1/2 | 1/2 |
| Network Connectivity | Unconstrained | Unconstrained | Constrained | Constrained | Constrained |
| Weight Symmetry | Not required | Required | Arises with learning | Not required | Arises with learning |
| Feedback Magnitude | Strong feedback | Weak nudging | Weak error | Weak nudging | Weak error |
| Error Encoding | Change in firing rate | Difference in neuronal activity | Activity of error neurons | Difference apical/somatic activity | Ratio bursts/single spikes |
| Update Rule | Temporal Hebbian | Contrastive Hebbian | Hebbian | Apical/somatic error | Burst rate |
| Update Timing | Continuous | Timed | Timed | Timed | Continuous |
| MNIST Performance | ~1.9% | ~2-3% | ~1.7-1.8% | ~2.0% | ~1.1% |
| Possible Link to Observations in Neuroscience | Apical inputs affect postsynaptic activity, increased feedback activity upon new stimuli | Varying neural responses for different behaviors | Increased neural activity upon new stimuli | Apical inputs affect activity and plasticity | Apical inputs affect burst rate and plasticity |
| Bioplausibility | +++ | + | ++ | ++ | +++ |

The relationship between temporal dynamics and bioplausible deep learning has been explored before. For instance, this was done by using subsequent frames, usually in an unsupervised or self-supervised setting (Illing et al., 2020; Lotter et al., 2020), and more generally in the so-called temporal error learning framework (Wittenberg and Wang, 2006). Our model applies a similar principle but with a supervised target and at the level of neuronal dynamics.

#### 3.1.4 Electrophysiological observations that agree with our model

In biological neural networks, LTP and LTD are one of the most prevalent forms of synaptic plasticity, and various studies have shown that LTP is induced when presynaptic spikes precede postsynaptic ones. In the case of multiple spike pairs, this is consistent with our model in that an increase in postsynaptic activity would lead to LTP and a decrease in LTD. Interestingly, recent work suggests that classical STDP-inducing protocols might fail under physiological extracellular calcium concentrations, suggesting that additional mechanisms might be required to act on the intracellular calcium levels (Inglebert et al., 2020; Larkum et al., 1999). In pyramidal neurons, intracellular calcium levels can be modulated by backpropagating action potential-evoked calcium (BAC) spikes that arise when apical inputs arrive shortly after basal inputs, resulting in action potential bursts (Larkum et al., 1999). Our model is consistent with this notion that delayed feedback into the apical dendrite drives plasticity while basal feedforward input does not. Future neuroscience experiments should explore if high calcium concentrations resulting from BAC spikes and bursts are indeed suitable to restore LTP and LTD induction when using a classical STDP protocol (Inglebert et al., 2020).

Finally, our model requires feedback that is specific to every neuron. Therefore, the synaptic weights to the apical dendrite have very specific values that must be computed by some biological mechanism. In previous work, we showed that these weights can be learned in a bioplausible manner by an anti-Hebbian leaning rule Meulemans et al. (2020). In biology, anti-Hebbian learning rules appear in disinhibitory GABAergic synapses (Karri et al., 2007), suggesting that the target used for learning in our model would be fed back into excitatory neurons through disinhibitory circuits. This nicely relates our work to the role of coupled apical and basal inputs in learning and the regulation of this coupling by disinhibitory circuits (Williams and Holtmaat, 2019; Avital et al., 2019; Zhang et al., 2014), and therefore use connectivity that matches the requirements of our feedback-based target propagation framework. Future theoretical investigations should continue this line of work by looking beyond Hebbian-like learning rules and integrating the knowledge of BAC-firing dynamics, the effects of calcium on plasticity, and the role of disinhibitory circuits in bioplausible models of deep learning.

### 3.2 Limitations and future work

#### 3.2.1 Limitations

One property of our framework is that during learning it requires a top-down controller to continuously compare the actual to the desired network output while sending feedback to the lower hierarchies. Although such a feedback controller can be easily realized as a neural circuit (Meulemans et al., 2021a), it is not clear yet if the brain employs any type of control circuit for learning. Future work could look at whether the apical inputs going through disinhibitory circuits correspond to feedback inputs that drive neurons to the desired target activity.

Lastly, in our model, weights from the same neuron can be positive and negative or even transition from negative to positive and vice-versa, which is in conflict with Dale’s law. This is a common simplification of ANN models (Cornford et al., 2020). Violating Dale’s law, however, can be corrected using a bias in the postsynaptic activity to turn negative weights into weak positive weights (Kriegeskorte and Golan, 2019). Moreover, recent studies showed that with certain network architectures and priors, Dale’s law can be easily preserved while maintaining the same functional network properties (Cornford et al., 2020).

#### 3.2.2 Future work

After learning, our model predicts an expedited onset of pyramidal neuron activity upon feedforward input (Fig. 3) that is inversely correlated with the top-down feedback to alter neuronal activity. A related cortical micro-circuit hypothesis is that local inhibitory microcircuits projecting onto apical dendrites control the neuron’s excitability and that their control strength reduces during learning. In an experimental setting, this temporal shift as well as the feedback strength attenuation could be tested using simultaneous in vivo 2-photon calcium imaging of excitatory and inhibitory populations (as in (Han et al., 2019)) combined with a plasticity-inducing whisker stimulation paradigm.

At the computational level, follow-up studies should go beyond modeling phenomenological learning rules such as STDP into hierarchical networks. For example, one direction could be to develop a more detailed mechanistic sub-cellular model that accounts for the coupling of intracellular voltage and calcium dynamics that are being differently modulated by inputs to apical and somatic synapses. Such sub-cellular mechanistic models might also include multiplicative effects of the apical input (Larkum et al., 2004) as well as apical-induced bursting (Segal, 2018) to further close the gap between the correlation-based models used in computational neuroscience and experimental observations showing, for example, the diverse intracellular effects of calcium on learning and neuronal activity (Larkum et al., 1999, 2007; Larkum, 2013). Another logical future step would be to develop more explicit theoretical links between PC and our temporal Hebbian framework. This would require applying it to other problems, such as detecting deviations from learned time series (Garrido et al., 2009) or unsupervised image representations (Rao and Ballard, 1999), and comparing the reduction of feedback with the minimization of prediction errors or free energy (Friston and Kiebel, 2009). Showing such conceptual links would pave the way to design more cortical-like circuits that explain the features of predictive coding but avoid the problems emerging from explicit error neurons (Kogo and Trengove, 2015).

Another limitation of our work is that we use DH instead of STDP to train deep networks. This is due to the randomness induced by our implementation of spiking neurons using a Poisson model, which implicitly imposes noisy learning updates. Further work could use leaky integrate-and-fire neurons, which can reduce the effects of randomness and maybe integrate them with models that consider more detailed physiological phenomena such as calcium spikes. This would require computing feedback in an event-based network, which is a currently active area of research.

Finally, our framework can be leveraged for applications beyond neuroscience. The simplicity and locality of neuron learning makes it well-suited for event-based neuromorphic processors and on-chip learning applications. This would require integrating a simple PI controller in a neuromorphic processor and further theoretical work on implementing our learning set-up with leaky integrate-and-fire neurons. Given that STDP can induce energy-efficient representations (Vilimelis Aceituno et al., 2020), it is likely that training with STDP might even further improve the energy efficiency of neuromorphic devices. In addition, the fact that our framework can learn all weights in an online manner (Meulemans et al., 2021b) implies that a perfect model of the processor architecture is not required, which is a key problem when training neuromorphic devices off-line due to the so-called device mismatch (Pelgrom et al., 1989; Binas et al., 2016).

## 4 CONCLUSIONS

We present a new hierarchical learning framework in which the temporal order of neuronal signals is leveraged to encode a top-down error signal. This reformulation of the error allows us to avoid unobserved learning rules while at the same time being consistent with classical ideas of predictive coding. Our work is a crucial step towards a more detailed understanding of how temporal Hebbian and STDP learning can be used for supervised learning in multilayer neural networks.

## AUTHOR CONTRIBUTIONS

P.V.A, M.T.F., and B.F.G designed the project and the experiments. P.V.A developed the mathematical STDP framework. P.V.A performed the single neuron learning simulations. M.T.F. performed the network simulations. P.V.A, M.T.F., and B.F.G wrote the paper. R.L. provided biology insights about the project on both theoretical and experimental neuroscience and wrote part of the discussion section.

## ACKNOWLEDGMENTS

We thank Christoph van der Malsburg, Alexander Meulemans, and Matthew Cook for fruitful discussions. We thank Maria Cervera for initially exploring the predictive coding properties of the standard DFC framework. We thank Sander de Haan for being part of the discussions and providing feedback on the paper.

## CODE AVAILABILITY STATEMENT

We will upload the code on the ETH GitLab repository upon acceptance and include a public download link in the paper.

## APPENDIX

### 1 Cost functions for Differential Hebbian in single layer networks

In this section, we show that the DH rule minimizes the cost functions mentioned in the main text. Even though we are referring to the simple neuron example represented in Figure 2, the results naturally hold for any single-layer linear classification setting.

I.1 MSE loss as a cost function

For the MSE loss cost function, ℒ, we note that the presynaptic activity is fixed, which gives us the update rule

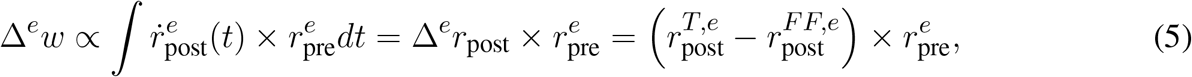

where *w* = (*w*_*A*_, *w*_*B*_), Δ*w* = (Δ*w*_*A*_, Δ*w*_*B*_), 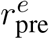 is the presynaptic activity vector, and *e* corresponds to the training example. For that specific example, the loss can then be represented as

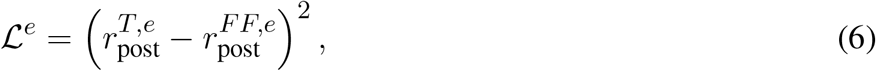

and using a standard derivative on the weights *w*_*A*_, *w*_*B*_ we recover the update rule in Eq. 5,

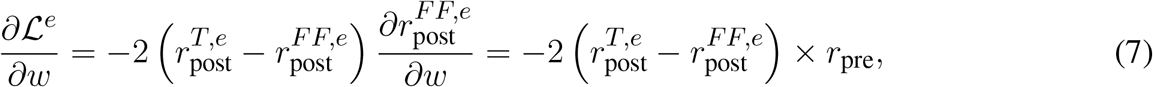

which corresponds to a standard gradient descent update.

To show that the feedback, ℋ, and time to target, 𝒯, are also minimized, we define 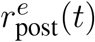 as a dynamical system whose attractive fixed point is the target activations for the different labels. Each example provides one initial condition of the dynamical system and the feedback ensures that the dynamics converge to the desired target.

#### I.2 Feedback strength as a cost function

In order to make the transition to the feedback strength, *H*, we note that

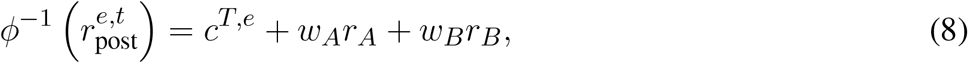

where 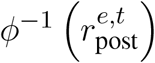 is the required membrane potential for the postsynaptic neuron to reach its target, hence, it is fixed while the right-hand side changes by learning. This allows us to rewrite the feedback strength cost function as

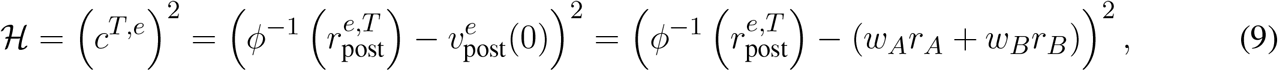

and, thus, we can write the gradient of the feedback strength cost function as

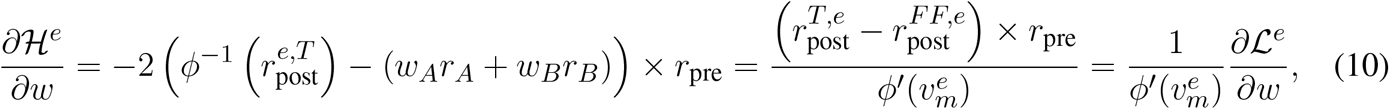

where the factor 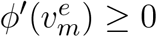 with 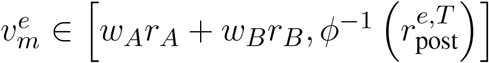 comes from applying the mean value theorem to the firing rate as a (positive definite) function of the membrane potential. Note that this is a very similar statement to the one presented in (Meulemans et al., 2021a).

#### I.3 Time-to-target as a cost function

In order to prove that the time delay cost function 𝒯 is minimized, we need to make assumptions on the feedback. Specifically, we will assume that the calculation of the feedback works directly on 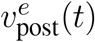, forcing it to reach its target value 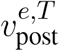. We will also assume that the feedback is computed by a PI controller, which stably converges to its target.

Since feedback pushes the firing rate to a target value, the dynamics of the membrane potential 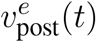 converge to a stable fixed point 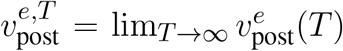. In order to avoid the limit *T* → ∞, we will consider the state space as a ball of radius *ϵ* centered around the target, and that the network has converged when it reaches that ball.

In a linear stable dynamical system, the activity evolves according to

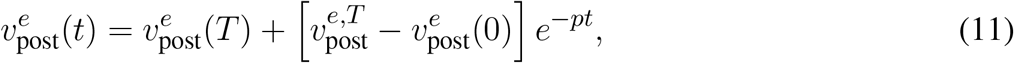

where *p* is the rate of convergence given by the projection of 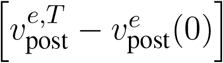 on the eigenspace of the feedback, which is positive as long as the feedback is stable.

Given the activity evolution, we can compute the time it would take for a training example to reach the target, *τ*^*ϵ*^, by

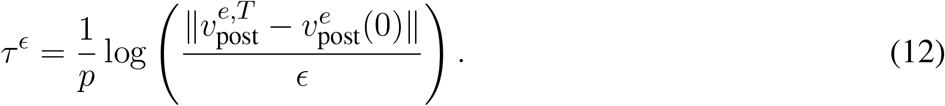

We note that if the system is nonlinear but monotonic, as it would be if the feedback is computed using the firing rate 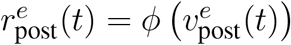, the previous formula would not work. Then, we would have to consider the projection 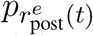 as a state-dependent exponent. While this would not have a closed-form solution, it can be bounded through the Lyapunov exponents of the network.

### II Extension to multilayer networks

In this section, we argue that the DH learning rule allows learning in multilayer networks for all the loss functions discussed in the previous section.

The gist of our argument is that the feedback defines a target activity for all layers of the network. Each layer has its own cost function, and the global cost function of the network is a positive semi-definite composition of the layerwise cost functions. Hence, global optimization is guaranteed by learning in each layer.

#### II.1 Differential Hebbian has the same fixed points as the dendritic-error learning rule

In this section, we show that given a feedback signal that successfully pushes the whole network to its target activities 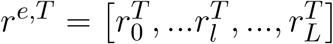, the DH learning has *r*^*e,T*^ as the fixed point, so

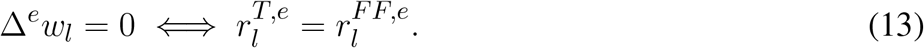

We start with the first layer. Given that the input is static and has a predefined target activity 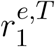, then

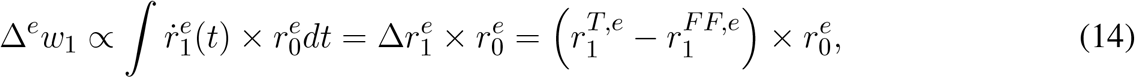

which has a single attractive fixed point for the learning dynamics at 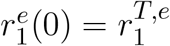.

Now, we can proceed by induction. For any layer *l* + 1, the DH learning rule will give the following weight update

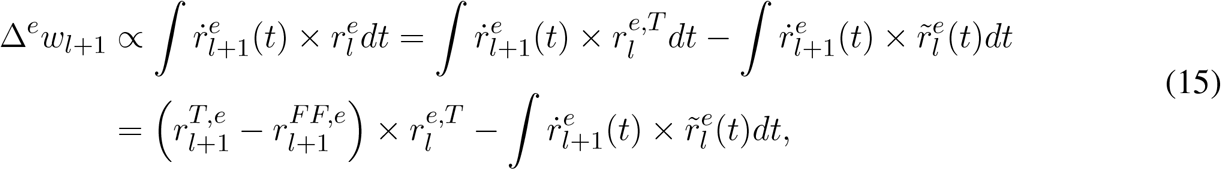

where 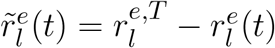 is the difference between the firing rate in layer *l* at time *t* and its target.

After enough learning, we have that 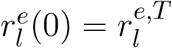. This implies that there is some epoch after which the learning rule in Eq. 15 becomes

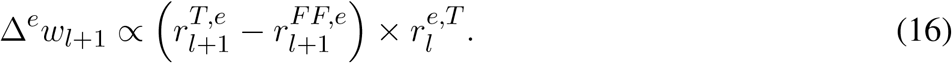

Which will eventually converge, giving us 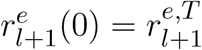. This can be iterated for all layers, following that a fixed point of dendritic-error learning from Eq.16 is also a fixed point of DH learning.

Note that the calculations presented are based on a single example, so the fixed point relates to a single class or label. Having an underparametrized model implies that there will be trade-offs such as the fixed point of the network parameters will not reaching zero loss for every example. How the loss for different examples is combined into a global loss in underparametrized systems is unknown in standard BP or DFC (Meulemans et al., 2020), as it is in our set-up.

#### II.2 Global and layerwise loss functions in deep networks

In this section, we define a loss function for every layer of the neural network and show that the global cost function is a positive definite composition of those rules.

We consider the cost functions for a given layer *l* denoted as ℒ_*l*_, ℋ_*l*_, 𝒯_*l*_. All of those will be defined on one layer assuming that the input to that layer is fixed to its target activity. We will also assume that the feedback is given by an oracle that knows from the beginning what the target activity of each neuron is and exactly the required feedback for the neuron to remain fixed to its target when the rest of the network is at equilibrium. Although this might be practically unfeasible in nonlinear systems, in practice, the appropriate PI controllers can provide good approximations.

First, we start with the ℒ global cost function. It is clear that the error is given by the last layer, so ℒ = ℒ _*L*_. Second, we consider the ℋ global cost function. The total amount of feedback that is given to the network is the sum of the amount of feedback given to each layer, so ℋ = ∑ _*l*_ ℋ _*L*_. Finally, for the time-to-target, 𝒯, the final layer can reach an equilib rium only if al l the layers have reached theirs. Hence, this takes as long as the slowest layer, so 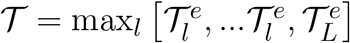.

From Section I, we know that reaching the fixed point of DH learning minimizes the local cost functions ℒ_*l*_, ℋ_*l*_, 𝒯_*l*_ if the presynaptic activity is static. By applying the inductive argument from Section 2.2, we know that throughout the learning the presynaptic activities will be approximately static, so the layerwise loss functions will be minimized. Finally, since the global losses are a positive definite combination of the layerwise losses, minimizing the local losses also leads to a minimization of the local ones.

### III STDP as a noisy version of Differential Hebbian

Although DH is supposedly equivalent to STDP with the right weight update kernel (Zappacosta et al., 2018), we found in Figure 4 that the weight updates with STDP are very noisy. We argue that the reason for this is that the conversion from rates – which we use to compute the appropriate feedback – to spike trains – which are required for STDP – induces randomness due to the use of a Poisson neuron model. A Poisson neuron model is by nature random, and we would expect some level of noise to be present all throughout the training. However, while the noise might remain at a given level, the learning signal does not; as the network is trained, the neurons start very close to their final targets, and therefore the change in neuronal activity – the learning signal – becomes much smaller. Maintaining a fixed level of noise but decreasing the signal strength implies that the learning signal eventually gets drowned in noise and cannot reach the level of precision necessary for competitive deep learning performances.

To test this argument we note that if every conversion from firing rates to spikes is noisy, we can reduce the noise by averaging for many conversions. Thus, we run our deep network for each epoch, but for every input, we perform multiple conversions of spike trains to firing rates and then average over the resulting STDP weight updates. We see that averaging indeed increases the performance achieved by the network (see Fig. S1). However, there is a limit to the feasibility of the approach, in that the number of averages required grows exponentially, thus scaling it up to reach state-of-the-art machine learning performance is not feasible.

**Figure S1.**
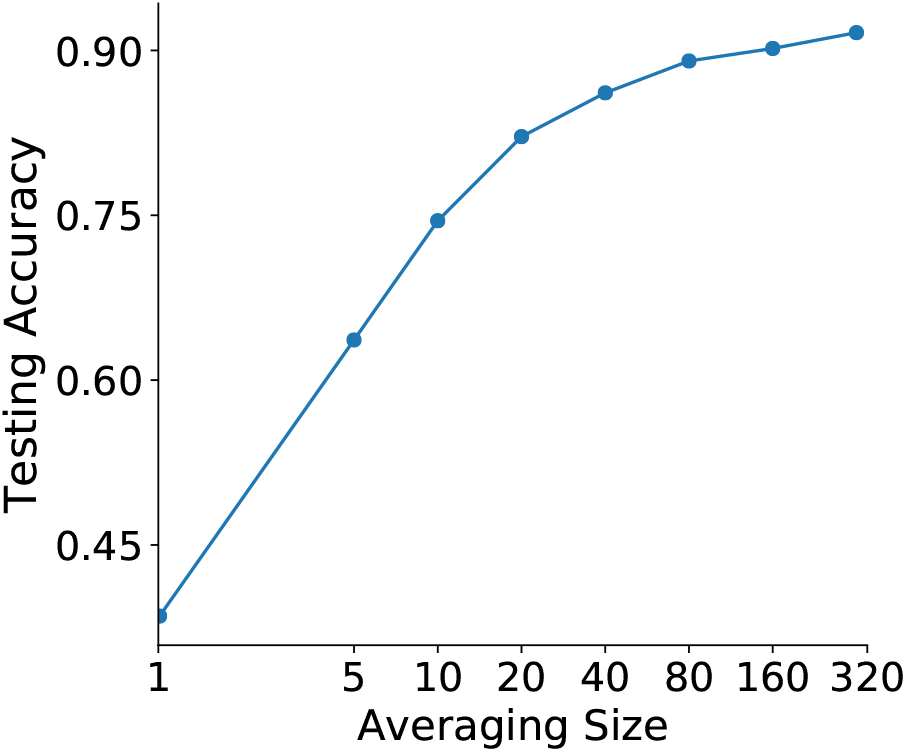
Averaging STDP increases its performance. We train a network with size 256×256×256 for the MNIST task but instead of using the DH learning rule we convert the rates to spikes with a Poisson neuron model and u se the STDP learning rule. We observe a steady increase in performance with the number of conversion samples.

### IV Simulations

#### IV.1 Description of the hyperparameter searches

The set of hyperparameters chosen for reporting values was selected by the best validation accuracy of all the training epochs, using 5000 validation samples extracted from the training set. We use the Tree of Parzen Estimators hyperparameter optimization algorithm (Bergstra et al., 2011) based on the Hyperopt (Bergstra et al., 2013) and Ray Tune (Liaw et al., 2018) Python libraries.

#### IV.2 Description of training

##### Weights and neurons initializations

The feedforward network weights are initialized with the Glorot-Bengio normal initialization (Glorot and Bengio, 2010). Unlike in (Meulemans et al., 2021a,b), the feedback weights here are kept frozen throughout the training and initialized to pre-trained values.

##### Activation functions

For the hidden neurons we use the sigmoid activation and for the output neurons we use a linear activation with a softmax readout.

##### Optimizer

To perform the reported MNIST experiments, we use Adam optimizer (Kingma and Ba, 2014) for the forward weights, which improves results compared to vanilla SGD. As mentioned in Meulemans et al. (2021a), Adam was designed for BP updates, so it is most likely not an optimal optimizer for DFC, which uses MN updates.

##### Differential Hebbian updates computation

We compute the DH updates throughout the time series of the pre- and post-synaptic firing rates. We compute the postsynaptic change in activity at each timestep and then multiply the result by the corresponding presynaptic activity,

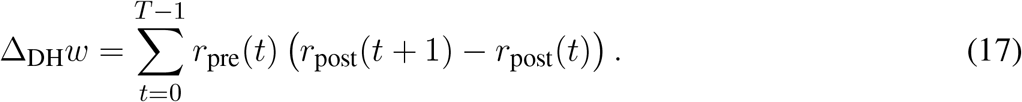

##### STDP updates computation

We compute the STDP updates by considering the time series of the pre- and post-synaptic rates. First, we convert the rates into spike trains by sampling from a random uniform distribution in the interval [0, 1] and comparing it with the firing rate at each timestep. Then, we convolve the presynaptic spike train with the following STDP kernel

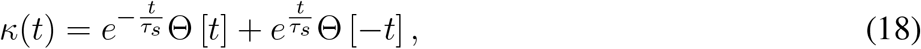

where Θ [*t*] is the step function and we set *τ*_*s*_ = 9.5 so that the decay rate is 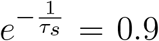 (for Fig. 2). Finally, we compute the weight update as the dot product of the convolved presynaptic spike train and the postsynaptic spike train,

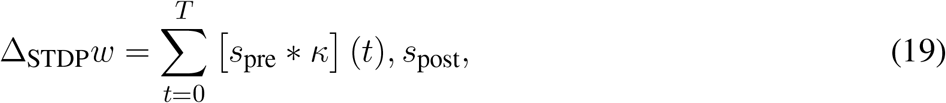

where *s*_pre/post_(*t*) is a spike train encoded as a vector of zeros and ones and * is the convolution operator.

#### IV.3 Description of the measures

Here, we describe how the measures reported in Figure 3 were obtained:

- **Feedback:** the feedback is measured as the amount of apical signal sent to each neuron, represented by *c*(*t*) Eq. 1. We report the average feedback strength per neuron across layers throughout the training epochs as the L2 norm (as in Meulemans et al. (2021b)).
- **MSE loss:** the mean squared error loss is computed taking the network’s output without feedback and the true labels throughout the training epochs.
- **Time-to-target:** the time-to-target was computed using DH learning with the original weak feedback setting of DFC (Meulemans et al., 2021a) as otherwise, the strong feedback setting (Meulemans et al., 2021b) immediately pushes the network’s output to the correct targets. For this, we measure the number of time steps taken to reach an *ϵ*-distance to the correct targets, where *ϵ* was taken as the maximum distance between the network’s output and the output targets for the last epoch.

#### IV.4 Resources and compute

To perform the reported MNIST experiments, we used GeForce RTX 2080 and GeForce RTX 3090 GPUs. We run for 40 training epochs and we did hyperparameter searches of 200 samples as in Meulemans et al. (2021a). The BP and DFC results are the same as reported in Meulemans et al. (2021b).

#### IV.5 Datasets and code licenses

For the experiments reported in this paper, we used the MNIST dataset (LeCun, 1998), with the license https://creativecommons.org/licenses/by-sa/3.0/.

For the implementation of the model, we used PyTorch (Paszke et al., 2019) and built upon the codebase of Meulemans et al. (2020, 2021a,b), which has the following licenses:

- Pytorch: https://github.com/pytorch/pytorch/blob/master/LICENSE
- Meulemans et al. (2020): https://www.apache.org/licenses/LICENSE-2.0

